# Molecular phylogeny of Catenulida (Platyhelminthes) with special focus on their diversity in Poland

**DOI:** 10.1101/2025.05.20.655055

**Authors:** Katarzyna Tratkiewicz, Jakub Baczyński, Ludwik Gąsiorowski

## Abstract

Catenulida is a clade of free-living flatworms found abundantly in freshwater habitats across the globe. Despite their ubiquitous distribution, catenulids remain poorly studied; most of the species are known only from the asexual forms that lack distinct, taxonomically useful characters. Accordingly, the studies of catenulid diversity require application of integrative methods that combine morphological and molecular data. Here, we report the survey of catenulid diversity in Central and Western Europe, with a special focus on the species found in Polish freshwaters. We collected and identified 13 distinct morphotypes that were subsequently sequenced for four molecular markers – *18S*, *28S*, *COI* and *ITS-5.8S*. The obtained sequences, together with reference data from other catenulid species, were used to infer the comprehensive phylogeny of the clade. The analysis revealed several well-supported clades within the largest catenulid family, Stenostomidae, highlighting the major challenges in catenulid taxonomy, such as unresolved species complexes of *Stenostomum leucops* and *Stenostomum simplex*. By tracing evolution of morphological, developmental and ecological characters on the phylogeny our study provides insight into major character transitions in the key lineages of catenulids.

## 1. Introduction

Catenulida is a small clade of microscopic turbellarians, that occupy sister position to all other flatworms (Egger et al. 2015; Larsson and Jondelius 2008; Laumer et al. 2015). Although they occur in both marine and freshwater habitats, most catenulid diversity is restricted to small and shallow water bodies with excessive organic matter such as eutrophic ponds, ditches, peatbogs, or rice fields (e.g., Diez and Schmidt-Rhaesa 2024; Larsson et al. 2008; Larsson and Willems 2010; Nuttycombe 1956; Nuttycombe and Waters 1938; Reyes et al. 2021; van der Land 1965; Yamazaki et al. 2012). Despite their prevalent occurrence, the diversity of catenulids remains understudied due to taxonomic challenges. First of all, these worms predominantly reproduce asexually and most species have never been observed in their sexual phase, limiting the taxonomic utility of reproductive characters – traits that are otherwise particularly important and reliable in the taxonomy of other flatworms (Brusa et al. 2020; Smith III et al. 2020). Second, certain catenulid species are known for their phenotypic plasticity (e.g., Nuttycombe 1956; Nuttycombe and Waters 1938; Rosa et al. 2015) resulting in the variability of some morphological character even within clonal animals with identical genetic background. Finally, many of the catenulid species are tiny (even for microturbellarian standards), relatively similar to each other, and difficult to anesthetize and fix, rendering their taxonomic identification problematic even for the trained experts (Sterrer and Rieger 1974). For these reasons, morphological investigations alone are not sufficient to reliably describe and quantify catenulid diversity, and the inclusion of molecular data is crucial for assessing their species richness.

While morphology-based investigations of catenulid faunas have been conducted for multiple world regions (e.g., Damborenea et al. 2011; Kolasa 1971; Kolasa and Young 1974a; Kolasa and Young 1974b; Marcus 1945; Noreña et al. 2005; Nuttycombe 1956; Nuttycombe and Waters 1938; Reyes et al. 2021; van der Land 1965), the systematic surveys of their diversity with the use of molecular markers are much more limited. The pioneering works of Larsson et al. focused on Swedish catenulids (Larsson et al. 2008; Larsson and Jondelius 2008; Larsson and Willems 2010) provided molecular tools and reference sequences for multiple species, enabling further molecular investigation of the catenulid diversity. The study was followed by systematic survey of the catenulid fauna in rice fields of Japan (Yamazaki et al. 2012), and recently in diverse aquatic habitats in Northern Germany (Diez and Schmidt-Rhaesa 2024). Extensive sequencing efforts have also been done in the symbiont-hosting genus *Paracatenula,* with sequences available for multiple species from Caribbean, Mediterranean and Red Seas, as well as Pacific Ocean (Gruber-Vodicka et al. 2011). Additional molecular data from single species have been also reported from Brazil (Rosa et al. 2017; Rosa et al. 2015), Thailand (Ngamniyom and Panyarachun 2016), Austria (Egger et al. 2017), and British Columbia (Van Steenkiste et al. 2023). However, many regions of the world remain unexplored in terms of catenulid molecular diversity.

In this study, we systematically sequenced four molecular markers (*18S*, *28S*, *COI* and *ITS-5.8S*) of several catenulid species found across Central and Western Europe. While our primary focus was on the fauna of northeastern Poland, we also included specimens from France, Germany, England, and Italy. Additionally, we inferred the phylogenetic placement of *Stenostomum brevipharyngium*, a recently emerging catenulid model (Gąsiorowski 2025; Gąsiorowski et al. 2025; Gąsiorowski et al. 2023). By combining publicly available data with sequences generated in this study, we reconstructed the most comprehensive molecular phylogeny of the clade to date. We used this phylogeny to trace evolution of selected morphological, developmental and ecological characters, as well as to revise certain taxonomical issues and highlight those that warrant further investigation.

## 2. Material and methods

### Animal collection and morphological observations

The animals were collected throughout various sampling sites in Poland, Germany, France, UK and Italy. In addition, we also used a laboratory strain of *Stenostomum brevipharyngium*, originating from unknown locality in the US, that remained in the laboratory cultures for the last 25 years (Gąsiorowski et al. 2023). Details on the sampling localities and identified species are provided in Tab. 1.

**Table 1.** Sampling localities and catenulid taxa identified.

| **Locality** | **Coordinates** | **Species** |
| --- | --- | --- |
| Poland: Choszczówka | 52.373870<br>20.957787 | *Stenostomum* cf. *simplex*,<br>*Catenula confusa* |
| Poland: Jezioro Śmierci | 53.653945<br>21.571690 | *Catenula* sp. |
| Poland: Mirosławiec | 53.347750<br>16.037877 | *Stenostomum handoelense*,<br>*Stenostomum* cf. *gigerium*,<br>*Stenostomum virginianum* |
| Poland: Woźnice | 53.843889<br>21.635000 | *Stenostomum saliens*,<br>*Stenostomum* cf. *simplex* |
| France: streambed of Massane river | 42.494306<br>03.028528 | *Stenostomum* cf. *leucops* |
| France: Paris | 48.843417<br>02.358667 | *Catenula* sp. |
| Germany: Göttingen | 51.538130<br>09.935559 | *Stenostomum saliens*,<br>*Catenula lemnae* |
| Italy: Fanano | 44.119860<br>10.803956 | *Stenostomum* cf. *leucops* |
| UK: Wellcome Trust Wetlands Nature Reserve | 52.073942<br>00.184608 | *Stenostomum constrictum* |
| USA | - | *Stenostomum brevipharyngium* |

The worms collected in the wild were sorted under dissecting microscope, and – when possible – cultured as isolines (originating asexually from a single individual). In successfully established isolines, some individuals were used for microscopic investigation, while others were used for the DNA extraction. If the worms were not reproducing asexually, the individuals used for the DNA extraction were documented directly.

For microscopic analysis, the worms were mounted in the drop of medium on microscopic slide, with the cover slip supported on the plasticine feet, and anesthetized using either low concentration of MgCl_2_ (1-1.4% w:v), cooling at 4°C for approximately 1 minute, or a combination of both. The anesthetized worms were imaged at the Nikon Eclipse NI-SSR with a Nikon DS-Ri2 camera. The microphotographs of the animals were adjusted for brightness and contrast in Fiji (Schindelin et al. 2012).

### DNA extraction, PCR amplification and sequencing

DNA was extracted using chelating agent – Chelex 100 resin (Bio-Rad, Hercules, Canada, USA), following procedure from Fells et al. (2023). Collected worms were transferred into appropriate clean culture medium or syringe-filtered ambient water and starved overnight. Single worms were collected into 1.5 ml Eppendorf plastic tube in a drop of medium and frozen at -70°C. Samples were then thawed and re-frozen repeatedly for total of three times. After the final thawing cycle, 100 μl of 10% Chelex water solution was added to each tube. Samples were incubated at 95°C for 30 minutes in thermomixer with interval mixing of 1000 rpm for 30 seconds every 2 minutes. Finally, the extracts were centrifuged at 10 000 rpm for 5 minutes and the supernatant with DNA was collected into clean 1.5 ml Eppendorf tube. PCR was performed with 2.5 μl of isolated DNA, 0.25 μM of forward and reverse primers and 12.5 μl of NZT Taq II 23 Green Master Mix (NzyTech, Lisboa, Portugal). The final volume of PCR mixture was 25 μl. Four genes were amplified – *18S rRNA*, *28S rRNA*, *COI* and *ITS-5.8S* with following pairs of primers: 4fb+1806R for *18S* of Stenostomidae, F19+R993(5R) for *18S* of Catenulidae, LSU5+L1642R for *28S*, COI5B+COI3B for *COI* and ITS5+ITS4 for *ITS* (Tab. 2). The PCR protocol consisted of 3 minutes of denaturation at 95°C, followed by 35 cycles comprising 30 seconds at 94°C, 30 seconds at 43–53°C and 30 seconds at 72°C. Final extension step was performed for 5 min at 72°C. PCR products were purified using Syngen PCR Mini Kit (Syngen Biotech, Wroclaw, Poland). The specificity of the PCR reaction was confirmed with gel electrophoresis. Purified DNA was sequenced using Sanger method by Genomed S.A. (Warsaw, Poland). Regarding their length, *18S* and *28S* genes were sequenced using both forward and reverse primers, and then assembled using Geneious Prime v. 2023.0.3. For *COI* and *ITS* genes respectively COI5B and ITS5 primers were used for sequencing. The sequences were manually trimmed for primers and ambiguous sites at the ends in Geneious Prime. The sequenced gene fragments were deposited in GenBank under accession numbers PQ887791–PQ887810, PQ895509–PQ895524, PV019496–PV019509 and PV403740–PV403751.

**Table 2.**
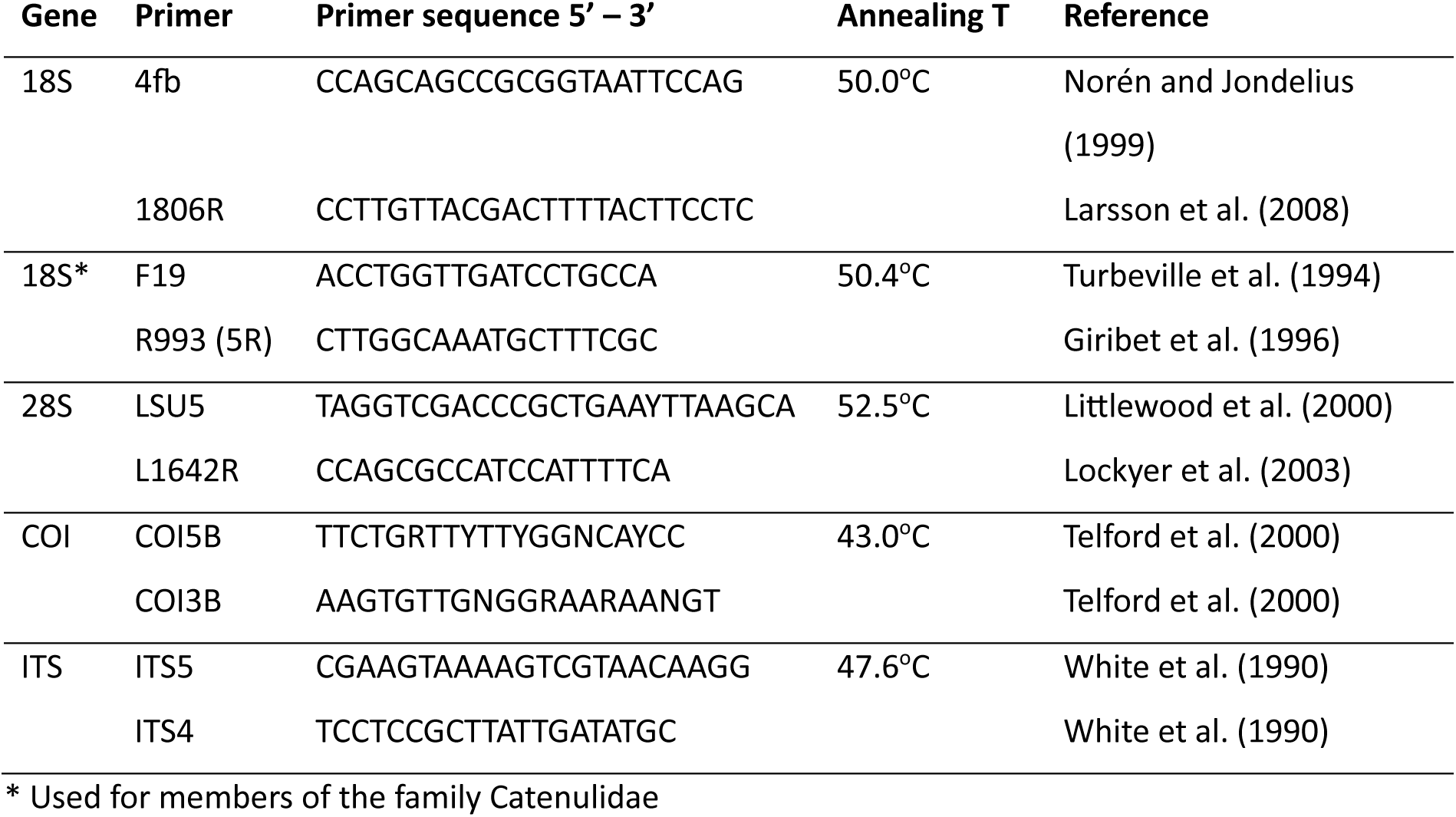
Primers used for PCR amplification and sequencing.

### Phylogenetic analyses

We assembled set of reference sequences of catenulids and other flatworms from GenBank (Tab. S1). Sequences from *18S*, *28S*, and *ITS* regions were aligned using MAFFT v7.271 (Katoh and Standley 2013) with automatic algorithm selection (--auto). For *COI*, a codon-aware alignment was generated using MACSE v2 (Ranwez et al. 2018). Ambiguously aligned regions were removed using the ’automated1’ trimming algorithm in trimAl v1.2 (Capella-Gutiérrez et al. 2009). The best substitution model for each marker was fitted using ModelTest-NG v0.1.7 (Darriba et al. 2020): TIM2+I+G for *18S*, GTR+I+G for *28S*, and TVM+G for *COI* and *ITS-5.8S*. The final concatenated alignment comprised 3,844 nucleotide positions, representing 125 distinct terminals (File S1), and was divided into four marker-specific partitions with the appropriate substitution models. The maximum likelihood phylogeny was inferred with these partitions using IQ-TREE v. 1.6.12 (Nguyen et al. 2014). Additionally, separate approximate-likelihood phylogenies for each molecular marker were analyzed with FastTree (v 2.1.11) (Price et al. 2010) implemented in Geneious Prime using default parameters.

### Ancestral state reconstructions

We assembled character matrix that covers all species from our phylogenetic analysis (23 species of *Stenostomum*, one species of *Rhynchoscolex*, three species of *Catenula*, three species of *Paracatenula*, one species of *Retronectes* and nine species belonging to the rhabditophoran outgroup). The matrix (File S2) includes information on 12 morphological characters that are important in catenulid taxonomy, two developmental characters related to asexual reproduction, one behavioral and one ecological character. We used our own morphological observation to score the character states for the species that we sequenced, while for the remaining terminals we used original species descriptions along with information from systematic surveys and reviews on catenulids (Diez and Schmidt-Rhaesa 2024; Larsson and Willems 2010; Nuttycombe 1956; Nuttycombe and Waters 1938; Yamazaki et al. 2012).

Ancestral state reconstruction was conducted on the best-scoring IQ-TREE topology, pruned to a species-level resolution with 40 terminal taxa, representing unique species. Analyses were carried out in R using the *corHMM* package (Boyko and Beaulieu 2021) under the marginal likelihood framework, with 20 random restarts to avoid local optima. For each trait, we tested two models of character evolution: ER (equal rates; a single rate parameter for all transitions) and ARD (all rates different; a distinct parameter for each possible transition). When the more complex ARD model did not result in a significantly better fit (ΔAICc < 2; see Tab. S2), the simpler ER model was used for subsequent reconstructions.

## 3. Results

### Morphological characteristics of the sampled animals

Our collection yielded 15 isolates, defined as unique combination of a single morphotype from a single location, representing 13 distinct morphotypes (Tab. 1). These included four morphotypes of *Catenula* and nine of *Stenostomum*. With the exception of two *Catenula* morphotypes that could not be imaged at a resolution satisfactory for species identification (and were therefore used only in the molecular analysis), all remaining morphotypes could be assigned to the described species:

*Catenula lemnae* Dugès, 1832

(Fig. 1A, B), sampling site: Göttingen, Germany

**Fig. 1.**
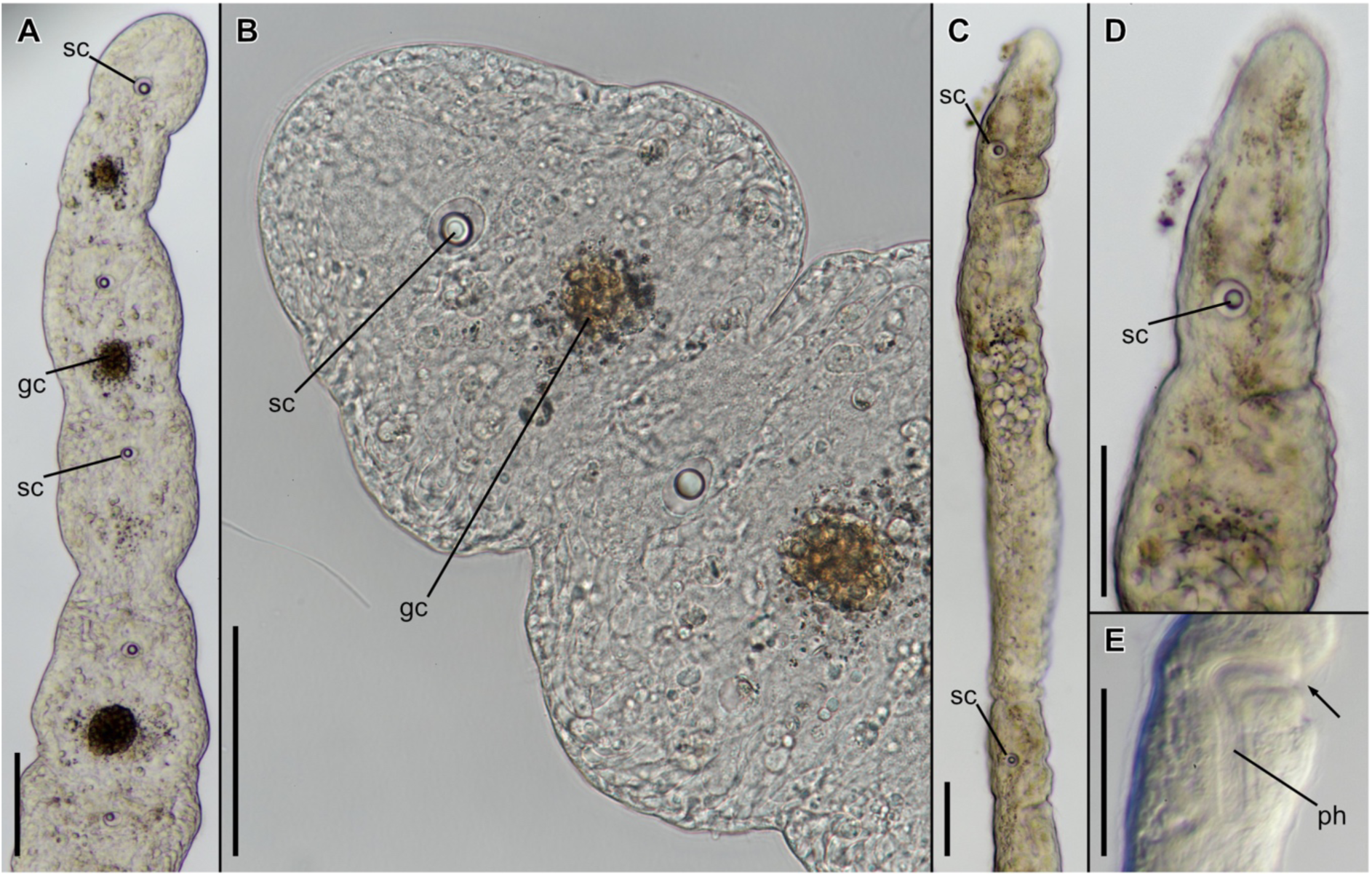
Morphology of *Catenula lemnae* (A, B) and *Catenula confusa* (C–E). Overviews of the fragments of zooid chains (A, C); magnified views of heads with visible statocysts (B, D) and pharynx (E). Abbreviations: gc, gut content; ph, pharynx; sc, statocyst; arrow, mouth opening. All scale bars equal 50 μm.

*Catenula* forming long chains composed of multiple short balustrade zooids (Fig. 1A), up to 20 zooids in one chain. Each zooid ca. 75-100 μm long with a well visible statocyst in the brain and short gut marked by dark content (Fig. 1B). Chains swim slowly but actively by undulating lateral movements of the entire conglomerate and can be often seen ascending into the water column. The animals can be cultured (although with difficulties) in the ambient water or MiEB12 medium with wheat infusion at 20°C.

*Catenula confusa* Nuttycombe, 1956

(Fig. 1C – E), sampling site: Choszczówka, Poland

*Catenula* forming short chains composed of two elongated and slender zooids (Fig. 1C). Each zooid ca. 450 μm long with a clearly visible statocyst in the brain (Fig. 1D). A rigid right-angled pharynx located behind the head connects ventral mouth opening with the gut (Fig. 1E). The animals swim fast with the ciliary action and bend only to change direction. The animals die out in captivity within few days, even when kept in ambient water.

*Stenostomum brevipharyngium* Kepner & Carter, 1931

(Fig. 2A – C), sampling site: unknown (laboratory cultures form the US)

**Fig. 2.**
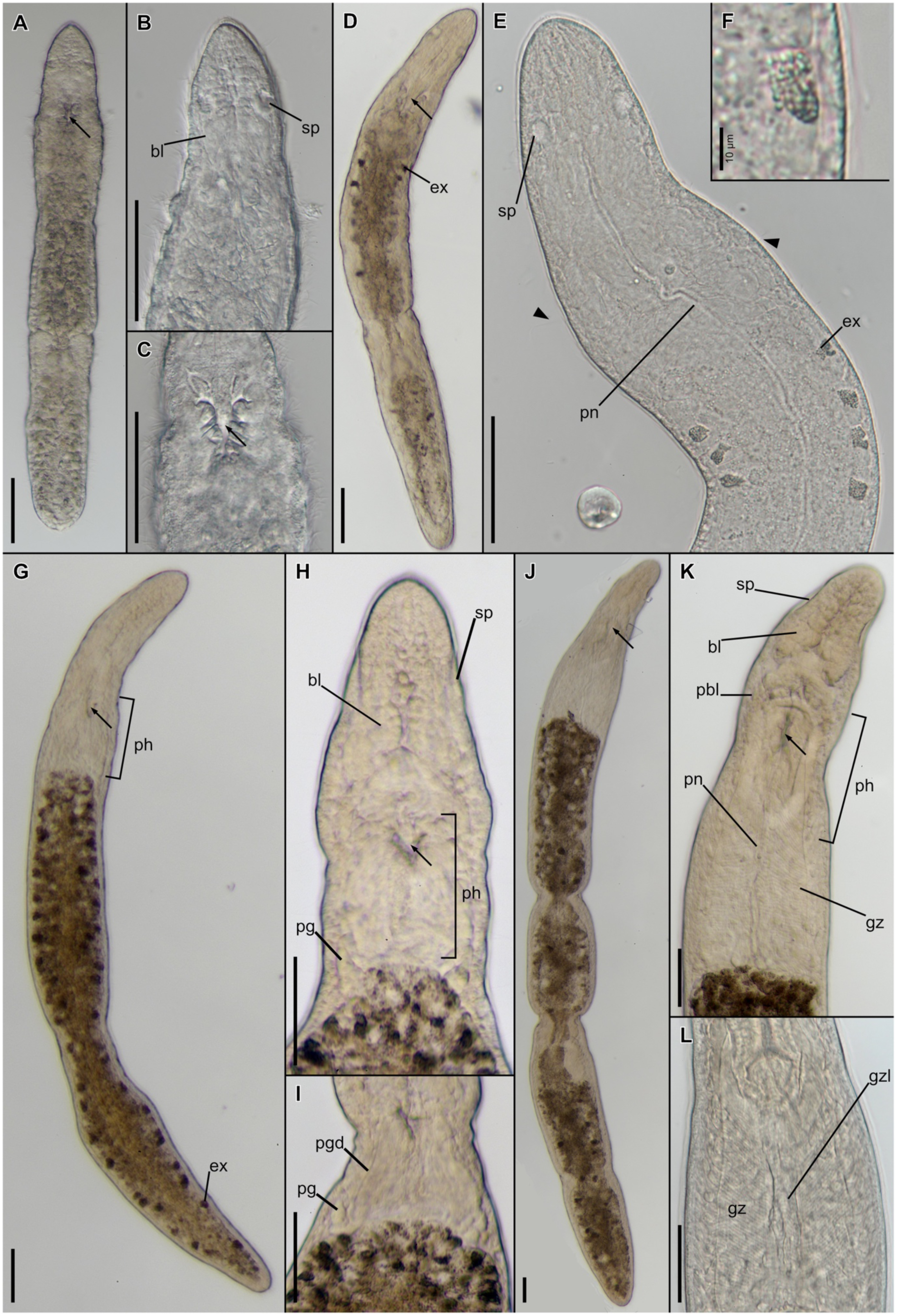
Morphology of *Stenostomum brevipharyngium* (A–C), *Stenostomum saliens* (D–F), *Stenostomum handoelense* (G–I), and *Stenostomum* cf. *gigerium* (J–L). Overviews of entire animals (A, D, G, J); magnified views of heads with sensory pits (B, E, H, K), mouth opening (C), excretophore (F), pharyngeal gland (I), and gizzard (L). Abbreviations: bl, brain lobe; ex, excretophores; gz, muscular gizzard; gzl, gizzard lumen; pg, pharyngeal gland; pgd, pharyngeal gland duct; ph, pharynx; pn, protonephridium; sp, sensory pit; arrow, mouth opening; arrowhead, long stiff cilia. Scale bars equal 50 μm unless described otherwise.

Small *Stenostomum* with single zooid reaching ca. 200-250 μm length (Fig. 2A). Body of equal width, with a slightly tapering anterior end and blunt posterior ends, gut reaches the most posterior extremity. Body covered in long, dense cilia of equal length (Fig. 2B and C). Inconspicuous brain lobes are visible in the head, accompanied by small, globular sensory pits that are almost fully enclosed in the head tissue (Fig. 2B). Ventrally located mouth is V-shaped and surrounded by clearly-visible lips, reinforced with rod-like elements (Fig. 2C). Mouth leads to a short pharynx, devoid of any easily discernable glands. Refracting bodies and excretophores are absent.

The worms can be easily cultured in Chalkley’s medium with wheat infusion at 20°C and fed with a cryptophyte *Chilomonas paramecium*. Under such conditions they reproduce asexually, but do not form chains of more than two zooids. The animals swim fast with the ciliary action, with their ventral side down.

*Stenostomum saliens* Kepner & Carter, 1931

(Fig. 2D – F), sampling sites: Göttingen, Germany and Woźnice, Poland

Small *Stenostomum* with single zooids reaching ca. 200-250 μm length (Fig. 2D). Body of equal width, with a slightly tapering anterior end and blunt posterior end. Gut extends to the most posterior extremity. An elongated protonephridium is clearly visible on the dorsal side of the animal (Fig. 2E). Elongated, stiff cilia distributed sparsely among shorter cilia across the entire body (arrowheads, Fig. 2E). Small, globular sensory pits are almost fully enclosed in the head tissue (Fig. 2E). Ventrally located mouth is V-shaped and surrounded by distinct lips (Fig. 2D). Mouth leads to a short pharynx, devoid of any discernable glands. A few excretophores are evident in the gut (Fig. 2D), composed of finely granular material (Fig. 2F). Refracting bodies absent.

The worms can be easily cultured in boiled tap water with wheat infusion at 20°C and fed with a cryptophyte *Chilomonas paramecium*. Under such conditions they reproduce asexually, but do not form chains of more than two zooids. The animals swim fast with the ciliary action, with their ventral side down.

*Stenostomum handoelense* Larsson & Willems, 2010

(Fig. 2G – I), sampling site: Mirosławiec, Poland

Large *Stenostomum*, with a single zooid reaching ca. 700 μm length (Fig. 2G). The head is long and slender, narrower than the trunk, both anterior and posterior ends are slightly tapering, gut reaches the most posterior extremity. Inconspicuous brain lobes are visible in the head, accompanied by elongated, shallow sensory pits that open widely to the environment (Fig. 2H). Small, ventral V-shaped mouth opening without lips (Fig. 2H) is located quite posteriorly in relation to the length of the entire head (Fig. 2G). Mouth leads to a short pharynx, with a pair of distinct pharyngeal glands (Fig. 2H and I). Each gland is composed of a glandular part, located at the border between pharynx and gut (Fig. 2H) and an elongated duct that extends along the pharynx (Fig. 2I). Numerous dark excretophores are present in the gut (Fig. 2G). Refracting bodies are absent.

As we found only one individual of this species, we did not attempt its husbandry. The animal was in non-reproductive stage and was actively swimming with the ciliary action with its ventral side facing down.

*Stenostomum cf. gigerium* Kepner & Carter, 1931

(Fig. 2J – L), sampling site: Mirosławiec, Poland

Large *Stenostomum*, with the largest recorded individual (with three fission planes) measuring ca. 1570 μm (Fig. 2J). The head is quite long and divided into narrower frontal part (ca. two fifths), and a broader posterior part that is as wide as the trunk. Thick, elongated protonephridium is evident on the dorsal side of the animal (Fig. 2J and K), and visible even under dissecting microscope, especially in the head. Dark gut reaches the most posterior extremity of the trunk. Several anatomical structure can be readily distinguished within the head. Distinct anterior brain lobes occupy the most anterior part of the rostrum and are accompanied by elongated, shallow sensory pits that open widely to the environment (Fig. 2K). Additionally, the elongated posterior brain lobes extend posteriorly along pharynx, reaching the level of mouth opening (Fig. 2K). Ventral mouth is slit-like (extending along A-P axis) and located at the border between narrow and broad parts of the head (Fig. 2J and K). It leads to a relatively short pharynx, which outline is visible as an oval of lighter coloration, occupying more or less 1/3 of the length of the head region (Fig. 2K). Behind the pharynx, there are two sets (left and right) of very prominent oblique muscle fibers that that form a muscular gizzard with a characteristic feather-like pattern (Fig. 2 K and L). The gizzard occupies entire space between pharynx and the gut, covering almost half of the head region. Fibers of the gizzard extend antero-laterally to the middle of the pharynx and end posteriorly to the border with the darkly colored gut. A narrow lumen of the gizzard is visible in the middle, between left and right sets of gizzard muscles, connecting posterior pharynx with the anterior edge of the gut (Fig. 2L). Refracting bodies and excretophores are absent.

The worms can survive for a couple of days in the filtered ambient water or Chalkley’s Medium, but they progressively become smaller and finally disintegrate, probably, due to starvation. So far, we did not succeed in attempts of culturing them. When found in the sample, the worms actively swam in the water column with the use of body ciliation.

*Stenostomum* cf. *leucops* (Dugès, 1828), morphotype 1

(Fig. 3A – D), sampling site: Massane’s streambed, France

**Fig. 3.**
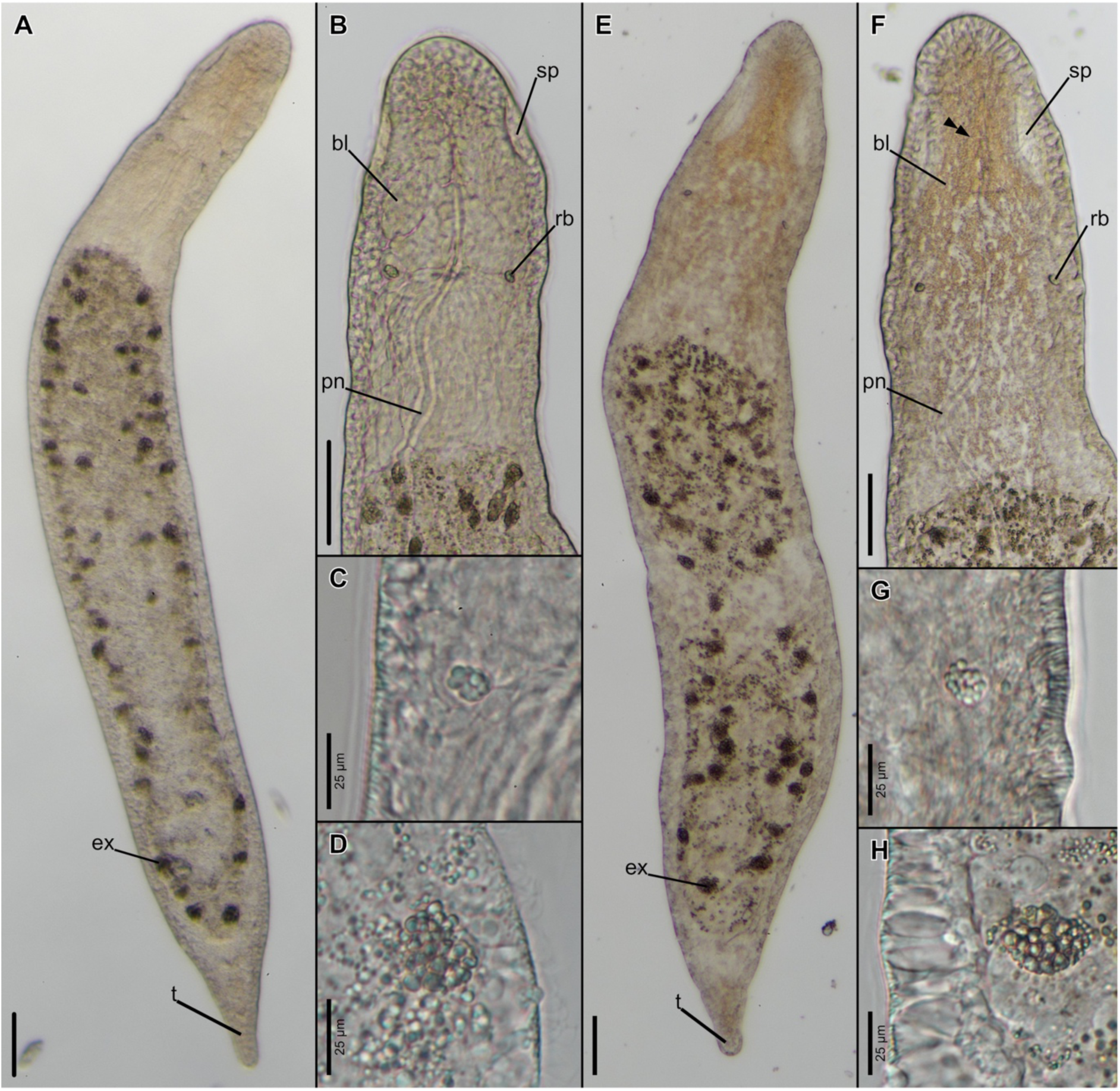
Morphology of *Stenostomum* cf. *leucops* – animals form France (A–D) and Italy (E–H). Overviews of entire animals (A, E); magnified views of heads with sensory pits (B, F), refracting bodies (C, G) and excretophores (D, H). Abbreviations: bl, brain lobe; ex, excretophores; pn, protonephridium; rb, refracting body; sp, sensory pit; t, tail; double arrowhead, skin pigmentation. Scale bars equal 50 μm unless described otherwise.

Large *Stenostomum*, with single zooids reaching ca. 1 mm length (Fig. 3A). The body is plump, and especially broad in the trunk region. The tip of the head is softly rounded, while posterior end of the body forms a distinct tail, devoid of gut tissue (Fig. 3A). A thick, elongated protonephridium is easily visible on the dorsal side of the animal (Fig. 3B). Distinct brain lobes occupy the anterior half of the head (Fig. 3B). Anteriorly they are accompanied by well-defined elongated, sensory pits that open widely to the environment, while posteriorly the brain lobes are equipped with the refracting bodies (Fig. 3B). Each refracting body is composed of smaller spherules (ca. 10 in the organ) that appear greenish in the transmitted light (Fig. 3C). Some of the gut cells are filled with excretophores (Fig. 3A), that are composed of numerous fine granules (Fig. 3D).

The worms can be easily cultured in boiled tap water with wheat infusion at 20°C with numerous unidentified single-cell eukaryotes thriving in the water. Under such conditions the worms reproduce asexually, but do not form chains of more than two zooids. The animals swim slowly with the ciliary action, with their ventral side facing down.

*Stenostomum cf. leucops* (Dugès, 1828), morphotype 2

(Fig. 3E – H), sampling site: Fanano, Italy

Large *Stenostomum*, with single zooids reaching ca. 1 mm length and general characteristic similar to the afore-described *S. leucops* (Fig. 3E). However, this morphotype can be distinguished from *S. leucops* morphotype 1 by presence of larger, more elongated sensory pits (Fig. 3F); yellowish pigmentation, especially prominent in the head, that is lacking from the area of sensory pits (Fig. 3F); and refracting bodies composed of more numerous (ca. 20) spherules (Fig. 3G). The gut of this morphotype contain excretophores, also composed of numerous fine granules (Fig. 3H).

The worms can be easily cultured in boiled tap water with wheat infusion at 20°C with numerous unidentified single-cell eukaryotes thriving in the water. Under such conditions the worms reproduce asexually, but do not form chains of more than two zooids. The animals swim slowly with the ciliary action, with their ventral side facing down.

*Stenostomum constrictum* Luther, 1960

(Fig. 4A – D), sampling site: Wellcome Trust Wetlands Nature Reserve, UK

**Fig. 4.**
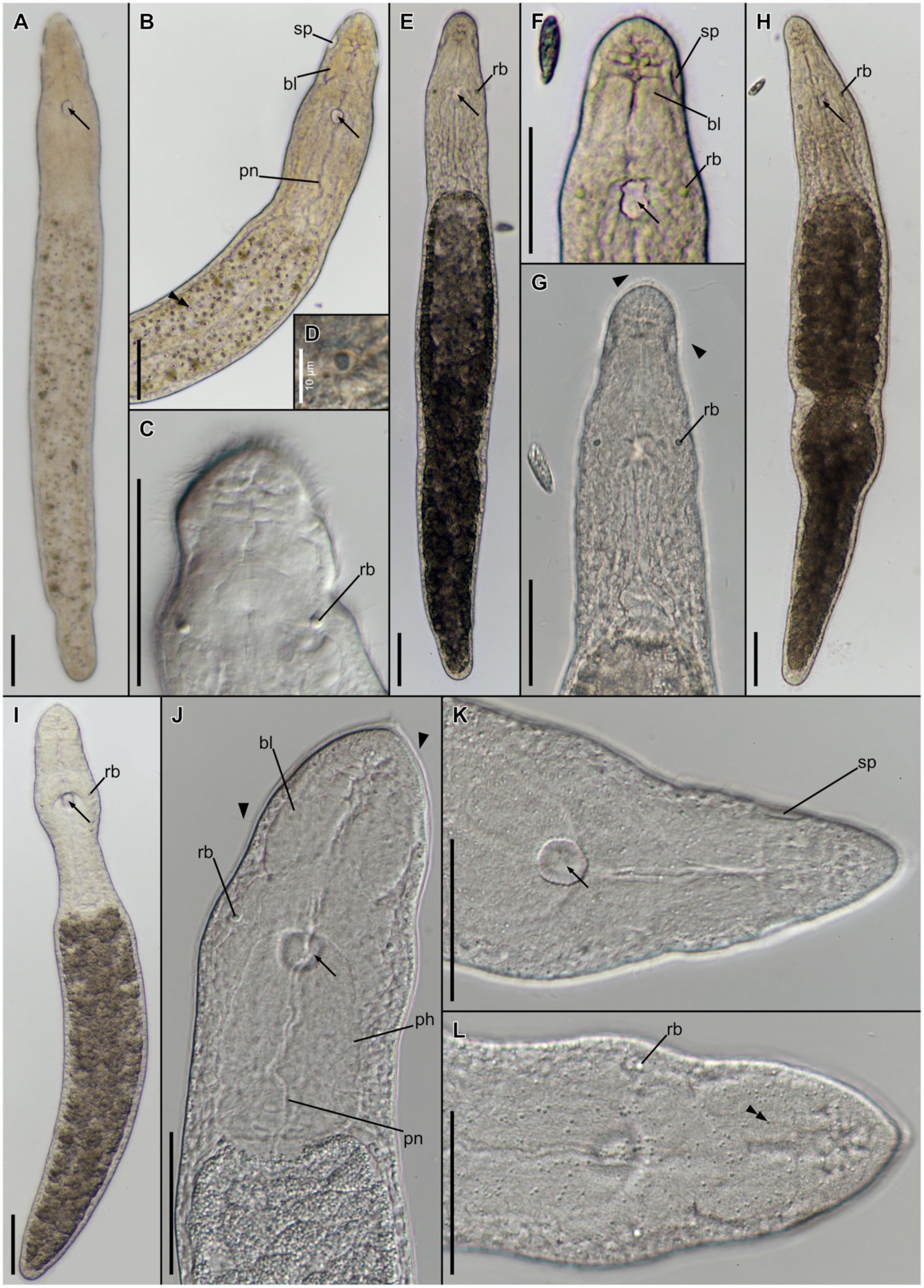
Morphology of *Stenostomum constrictum* (A–D), *Stenostomum* cf. *simplex* – specimen from Woźnice (E–G) and Choszczówka (H), and *Stenostomum virginianum* (I–L). Overviews of entire animals (A, E, H, I); magnified views of heads with visible mouth openings (B, F, K), heads with visible refracting bodies (C, G, J), refracting body (D), and skin spherules (L). Abbreviations: bl, brain lobe; pn, protonephridium; rb, refracting body; sp, sensory pit; arrow, mouth opening; arrowhead, long cilia; double arrowhead, skin spherules. Scale bars equal 50 μm unless described otherwise.

Large *Stenostomum*, with a single zooid reaching ca. 650 μm length (Fig. 4A). Body divided into head region, pharynx and trunk, with the head region being considerably narrower than the pharynx (Fig. 4B). Body tapers slightly posteriorly, however the gut reaches the most posterior extremity. The zigzagging protonephridium is visible on the dorsal side of the worm (Fig. 4B). Conspicuous brain lobes with the anterior metamery are visible in the head (Fig. 4B). Anteriorly the brain lobes are accompanied by small, elongated, sensory pits that open widely to the environment (Fig. 4B). Posteriorly the brain lobes are equipped with a pair of refracting bodies (Fig. 4C), each composed of single darker spherule enclosed inside a sack-like organ (Fig. 4D). Large, ventral, O-shaped mouth opening without lips is located at the anterior extremity of the pharynx (Fig. 4A and B). Pink-colored pigmented granules that are present inside skin cells give a spotty appearance to the worm (Fig. 4B). The remaining tissues, are generally less transparent than in other species investigated in this report, giving brownish-milky tint to the body of the worm and making visualization of the internal structures challenging. The excretophores are absent.

The worms can be easily cultured in Chalkley’s medium with wheat infusion at 20°C and fed with a cryptophyte *Chilomonas paramecium*. Under such conditions they reproduce asexually, but do not form chains of more than two zooids. The animals swim fast with the ciliary action, with their ventral side facing up and the mouth opening gaping widely.

*Stenostomum cf. simplex* Kepner & Carter, 1931

(Fig. 4E – H), sampling sites: Woźnice, Poland (Fig. 4 E – G) and Choszczówka, Poland (Fig. 4H)

Large *Stenostomum*, with a single zooid reaching ca. 620 μm length (Fig. 4E, H). Body of equal width, tapering slightly at both anterior and posterior extremities. The gut reaches the most posterior extremity of the body. Elongated, stiff cilia sparsely distributed among shorter cilia throughout entire body (arrowheads, Fig. 4G). Conspicuous brain lobes with the anterior metamery are visible in the head (Fig. 4F). Anteriorly, the brain lobes are accompanied by small elongated, sensory pits that open widely to the environment (Fig. 4F). Posteriorly the brain lobes are equipped with a pair of refracting bodies (Fig. 4E – H), each composed of single spherule, greenish in the transmitted light, that is enclosed inside a sack-like organ (Fig. 4G). Large, ventral mouth opening is round but slightly irregular, devoid of any lip-like structures (Fig. 4F). The excretophores are absent.

The worms can be easily cultured in Chalkley’s medium with wheat infusion at 20°C and fed with a cryptophyte *Chilomonas paramecium*. They also feed on *Vorticella*-like ciliates growing in the cultures. Under such conditions the worms reproduce asexually, but do not form chains of more than two zooids. The animals swim fast with the ciliary action, with their ventral side facing up and the mouth opening gaping widely.

*Stenostomum virginianum* Nuttycombe, 1931

(Fig. 4I – L), sampling site: Mirosławiec, Poland

Large *Stenostomum*, with a single zooid reaching ca. 500 μm length (Fig. 4I). Body of equal width, with the gut reaching the most posterior extremity. The head and pharynx region are very mobile, actively shifting their size and proportion due to frequent muscular movements. The zigzagging protonephridium is visible on the dorsal side of the worm (Fig. 4J). Small greenish spherules are distributed throughout the skin cells (Fig. 4L). Elongated, stiff cilia are sparsely distributed among shorter ones (arrowheads, Fig. 4J). Large, ovoid brain lobes are not distinctly divided into metamers (Fig. 4J). Anteriorly, the brain lobes are accompanied by small, elongated and very shallow sensory pits that are barely distinguishable in the light microscopy (Fig. 4K). Posteriorly the brain lobes are equipped with a pair of refracting bodies (Fig. 4J), each composed of single spherule, greenish in the transmitted light, that is enclosed inside a sack-like organ (Fig. 4J). Large, ventral mouth opening is regular, O-shaped and devoid of any lip-like structures (Fig. 4K). It leads to a stout, muscular pharynx, that is very clearly visible in the transmitted light (Fig. 4J). There are no distinct glandular structures associated with the pharynx. The excretophores are absent.

The worms can be easily cultured in Chalkley’s medium with wheat infusion at 20°C and fed with a cryptophyte *Chilomonas paramecium*. Under such conditions the worms reproduce asexually, but do not form chains of more than two zooids. The animals swim fast with the ciliary action, with their ventral side facing up and the mouth opening gaping widely.

### Molecular phylogeny of catenulids

Maximum likelihood analysis (Fig. 5) for the concatenated dataset recovered monophyletic Catenulida, divided into Stenostomidae and a clade uniting *Retronectes*, *Paracatenula* and *Catenula*, with the latter two forming sister groups. Within *Catenula*, we retrieved three well-supported clades, delimiting each of the sampled species: *Catenula turgida*, *Catenula macrura* and *Catenula lemnae*. The animal tentatively identified as *Catenula confusa* based on morphological studies was nested within *C. lemnae* clade. Within Stenostomidae, we retrieved four major clades with high bootstrap support: Clade 1 (*S. bryophilum* and *S. grabbskogense*), Clade 2 (*S. brevipharyngium*, *S. saliens*, *S. tuberculosum*, *S. gigerium*, *S. heebuktense*, *S. steveoi*, *S. handoelense*), Clade 3 (*S. lecuops* species complex, including *S. grande* and *S. sthenum*), and Clade 4 (*S. simplex* species complex, *S. sphagnetorum*, *S. constrictum*, *S. gotlandense*, *S. virginianum*, *Stenostomum* “island” group). Additionally, we recovered monophyletic *Rhynchoscolex simplex* and *S. arevaloi*.

**Fig. 5.**
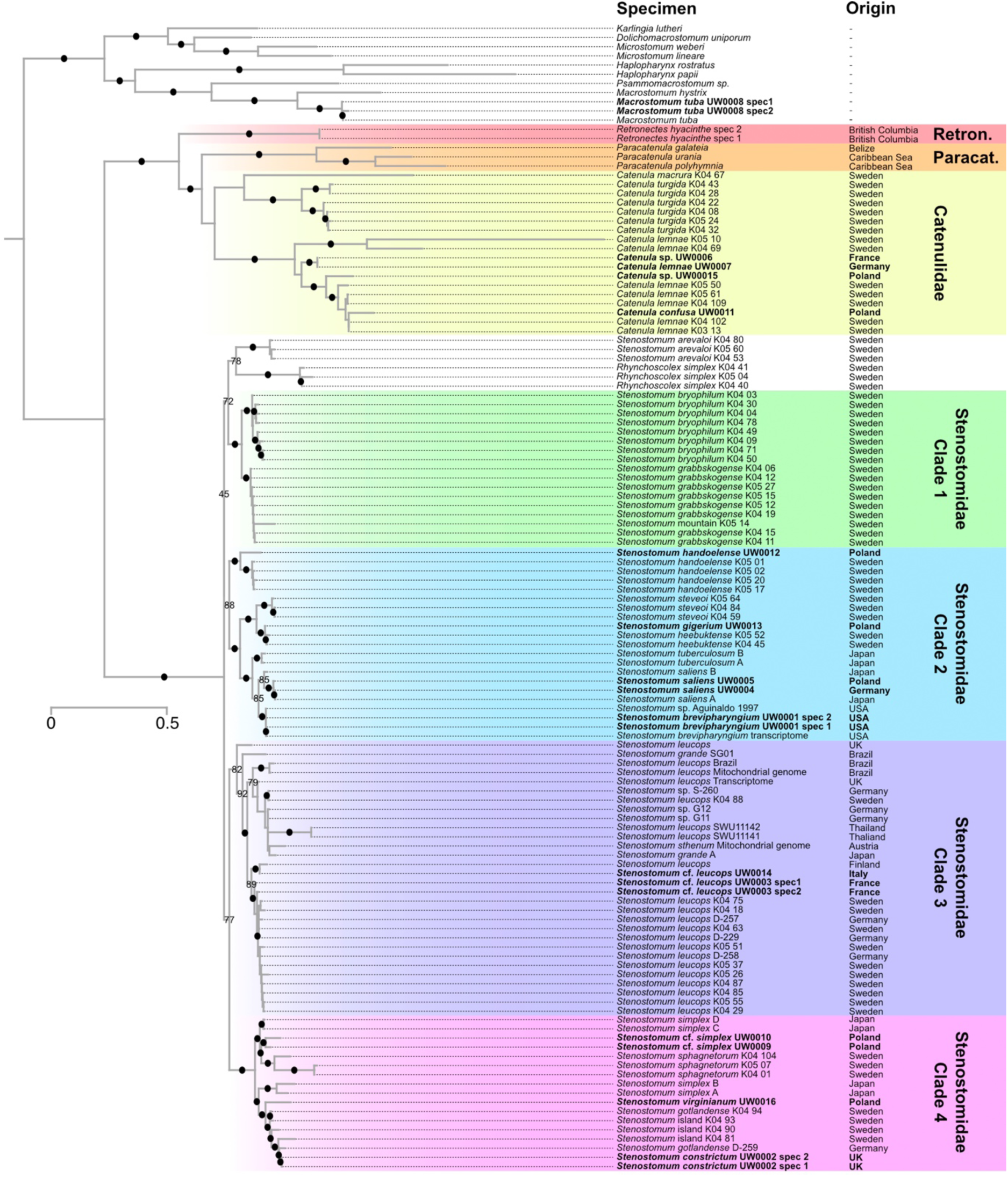
Molecular phylogeny of Catenulida inferred with a maximum likelihood approach from the concatenated *18S*, *28S*, *ITS-5.8S,* and *COI* datasets. Filled dots indicate bootstrap support >95, lower values are provided also for other important nodes. The original tree with bootstrap values for all nodes can be found in supplementary File S3. Terminals in bold indicate newly sequenced individuals. “Retron.” stands for Retronectidae and “Paracat.” For Paracatenulidae.

Our final analysis failed to recover monophyletic *Stenostomum*, as *R. simplex* grouped together with *S. arevaloi* within a clade encompassing Clades 1 and 2. The bootstrap support for this assemblage was, however, relatively low in comparison with other branches. The *S. lecuops* species complex can be subdivided into several well-supported and sometimes morphologically distinct clades, that likely represent cryptic species: a clade that includes animals from France, Germany and Sweden (corresponding to *S.* cf. *leucops* morphotype 1), a clade with animals from Finland and Italy (corresponding to *S.* cf. *leucops* morphotype 2), a clade with animals from Japan, Austria, Thailand, Germany, Sweden, Brazil and the UK, and two separated lineages with single individuals from Brazil and the UK. Importantly, the sequences of the specimens that have been identified as *S. grande* do not form a clade.

Finally, within Clade 4, we recovered division into two well-supported clades. The first one includes *S. sphagnetorum*, *S.* cf. *simplex* from Poland and some *S.* cf. *simplex* from Japan. The second clade contains *S. constrictum*, *Stenostomum* sp. (D-259) from Germany, *S. gotlandense*, *S. virginianum*, three undescribed *Stenostomum* species from Sweden (“island group”: undescribed species 1 – 3 from Larsson and Willems (2010)), and some *S.* cf. *simplex* from Japan.

### Ancestral state reconstruction

Reconstruction of the ancestral states for eight characters revealed phylogenetically informative distribution (Fig. 6). Regarding habitat, the last common ancestor of Catenulida occupied freshwater environments, with independent transitions to marine habitats in Retronectidae and Paracatenulidae. Paratomy was recovered as evolved convergently in *Microstomum* and catenulids and subsequently lost in *Retronectes* and *Paracatenula*.

**Fig. 6.**
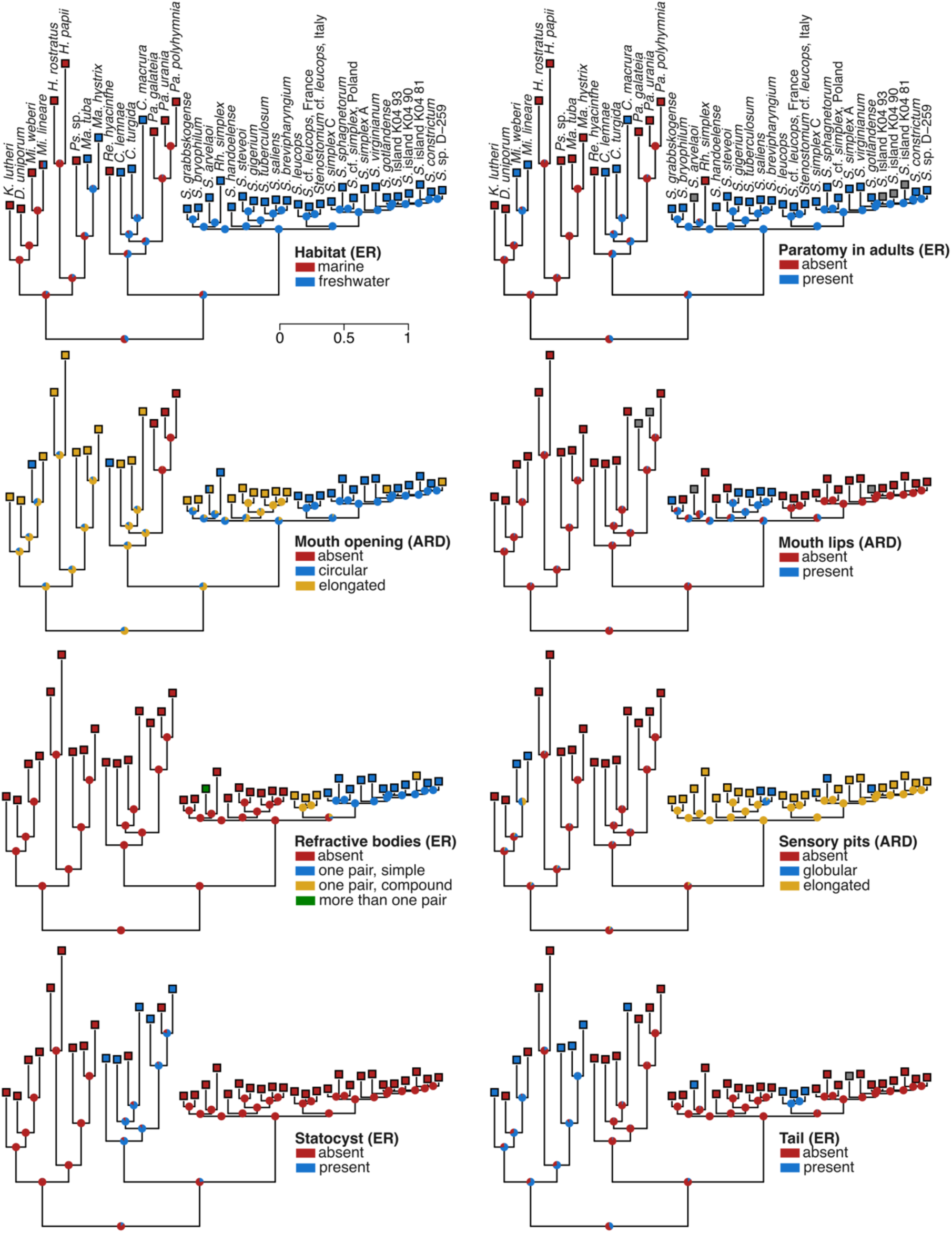
The ancestral state reconstruction of the ecological, developmental and morphological characters in catenulids. The tree topology was inferred based on a maximum-likelihood analysis shown in Figure 5. The pie charts depict the marginal likelihood of the character states at each node. For each reconstruction, the best-fitting model is indicated as either ER (Equal Rates) or ARD (All-Rates-Different). Grey squares indicate missing data.

The transition from elongated to circular mouth opening was reconstructed at the ancestral branch of Stenostomidae, with a reversal at the base of Clade 2. Mouth lips evolved in Stenostomidae, with a strong support for their presence in the common ancestors of Caldes 1 and 2.

Regarding sensory structures, refractive bodies were reconstructed as absent in the last common ancestor of both Catenulida and Stenostomidae, having evolved independently three times: as multiple pairs in *S. arvelaoi*, as a single pair of compound structures in Clade 3, and as a single pair of simple organs in Clade 4. In contrast, both sensory pits and statocysts evolved only once in catenulids; the former in the lineage of Stenostomidae, and the latter in the last common ancestor of Retronectidae, Paracatenulidae and Catenulidae.

Finally, the posterior tail evolved independently three times within Catenulida: in *C. macrura*, *S. arvelaoi* and *S. leucops* species complex. The remaining characters were not phylogenetically informative (Figure S1), although in some cases this was likely influenced by missing data in the matrix, which causes the reconstruction algorithm to assign equal likelihoods to all states. For instance, upside-down swimming appears restricted to Clade 4, but due to the data gaps for many of its members, the trait was reconstructed to have evolved multiple times independently within that clade (Figure S1).

## 4. Discussion

As our analysis presents the most comprehensive phylogeny of Catenulida published to date, we would like to use it as a basis for the brief discussion of catenulid taxonomy, including remarks on selected clades and species.

At the level of major clades, we retrieved the same topology as Van Steenkiste et al. (2023), supporting their finding of the paraphyly of Retronectidae (*sensu* Sterrer and Rieger (1974)), and a sister relationship between Paracatenulidae and Catenulidae. We also recovered a monophyletic Stenostomidae, consistent with multiple previous studies (Diez and Schmidt-Rhaesa 2024; Larsson et al. 2008; Van Steenkiste et al. 2023; Yamazaki et al. 2012).

Intriguingly, within Stenostomidae, we failed to recover *Rhynchoscolex* as a sister group to *Stenostomum*, a relationship robustly supported by several other studies (Diez and Schmidt-Rhaesa 2024; Larsson et al. 2008; Van Steenkiste et al. 2023). Instead, in our analysis *Rhynchoscolex simplex* was placed within *Stenostomum* (although with a relatively low support), as a sister species to *S. arevaloi*. Inspection of individual marker trees (Files S4–7), revealed that this topology is supported only by the *28S* dataset; in all other markers the species was recovered in the expected position as a sister taxon to *Stenostomum*. Given that *28S* is the longest gene in our dataset and contains the highest number of phylogenetically informative sites, the apparent inclusion of *R. simplex* within *Stenostomum* is likely an artifact. Sequencing additional species of *Rhynchoscolex* or additional markers for the species included in our analysis will be necessary do determine whether *Rhynchoscolex* indeed represents a derived lineage of *Stenostomum*.

Within Stenostomidae, we retrieved four well supported clades, united not only by molecular data but also by potential synapomorphies. For instance, species in Clade 2 share an elongated mouth opening with lips; those in Clade 3 are characterized by compound refractive bodies and the presence of a tail; and the species in Clade 4 possess simple refractive bodies and possibly exhibit upside-down swimming behavior. Considering that the genus *Stenostomum* comprises roughly half of all described catenulid species, its division into several molecularly well-supported and morphologically distinct genera might be useful in prospective taxonomical studies.

In future, a special attention should be devoted to resolving internal relationships of Clades 3 and 4. Clade 3 includes the *Stenostomum leucops* species complex, notorious for its poorly defined and paraphyletic species boundaries (Borkott 1970; Diez and Schmidt-Rhaesa 2024; Nuttycombe and Waters 1938; Rosa et al. 2015; Yamazaki et al. 2012). Here, we report a similar issue within Clade 4, which includes several yet undescribed species, collectively referred to as “island group” (Larsson et al. 2008; Larsson and Willems 2010). This clade includes *S. gotlandense* and morphotypes resembling *S. simplex* that likely represent a distinct species (Yamazaki et al. 2012, this study).

### *Taxonomic identity of* Stenostomum D-259 *and internal relationships within Stenostomidae Clade 4*

The animal recently reported by Diez and Schmidt-Rhaesa (2024) as *S. gotlandense* from Germany (individual *Stenostomum* D-259), recovered in Clade 4 in our analysis, also represents a separate, possibly undescribed species. This taxonomic confusion arises from the treatment of Diez and Schmidt-Rhaesa (2024), who considered all sequences labelled as *Stenostomum* “island” in Larsson et al. (2008) as *S. gotlandense*, although only the individual K04_94 actually represents this species. The three remaining individuals were originally reported as related, but morphologically distinct undescribed species (Larsson et al. 2008; Larsson and Willems 2010). The inclusion of additional sequences in our analysis shows that *Stenostomum* D-259 is more closely related to *S. constrictum* than to any other sequenced species, although the two are morphologically distinct. Nevertheless, regardless of the taxonomic status of *Stenostomum* D-259, both our study and that of Diez and Schmidt-Rhaesa (2024) demonstrate that the “*Stenostomum* island” clade is not restricted to the islands of Öland and Gotland, as originally suggested (Larsson et al. 2008), but also includes *Stenostomum* D-259 from Hamburg and *S. constrictum* that has been reported from Eastern Fennoscandia (Luther 1960), Eastern Europe (Kolasa 1973; Luther 1960) and Great Britain (this study).

Finally, two species belonging to Clade 4, *S. constrictum* and *S. sphagnetorum*, were historically treated as a single species – *S. unicolor* (Luther 1960). Although hardly distinguishable based on morphology, our analysis confirms that they represent two independent species that are in fact quite distantly related within Clade 4. Their apparent similarity likely reflects convergent evolution or independent retention of the plesiomorphic morphology.

### *Phylogenetic position of* Stenostomum brevipharyngium

One of our main goals was to reconstruct the phylogenetic position of *Stenostomum brevipharyngium*, a species that has recently emerged as a useful model for studies on various aspects of catenulid biology, with several established molecular techniques and resources (Gąsiorowski 2025; Gąsiorowski et al. 2025; Gąsiorowski et al. 2023). Our analysis indicates that *Stenostomum* sp. included by Aguinaldo et al. (1997) in their large scale metazoan phylogeny is conspecific with our cultures of *S. brevipharyngium*. We further demonstrate that *S. brevipharyngium* and the morphologically similar *S. saliens* are sister taxa within Clade 2, alongside *S. tuberculosum*, *S. gigerium*, *S. heebuktense*, *S. steveoi*, and *S. handoelense*. However, in contrast to the previous suggestion of Kolasa (1977), *S. brevipharyngium* and *S. saliens* are genetically well separated and morphologically different. For instance, *S. saliens* possesses excretophores and elongated, stiff cilia sparsely distributed on the body surface that are absent in *S. brevipharyngium*, in accordance with the original description by Kepner and Carter (1931). These findings support the treatment of both species as valid and distinct.

### *What is* Stenostomum gigerium?

A separate section of the taxonomic discussion must be devoted to the animal we provisionally identified as *Stenostomum* cf. *gigerium*. Originally described by Kepner and Carter (1931), this species was later synonymized with *S. tauricum* (Marcus 1945) by Nuttycombe and Waters (1938). Luther (1960) subsequently reassigned it to a new genus *Myostenostomum*, as *Myostenostomum tauricum*. Later, Rogozin (1992) redescribed the species as *Myostenostomum gigerium*. Further complications arose through taxonomic treatment of Noreña et al. (2005), who transferred the species to the now obsolete genus *Anokkostenostomum*. Ultimately, both *Myostenostomum* and *Anokkostenostomum* were transferred to *Stenostomum* based on morphological and molecular phylogenetic analyses (Damborenea et al. 2011; Larsson et al. 2008). Due to this convoluted taxonomic history, the species appears in the literature under multiple names and combinations.

The species we collected near Mirosławiec, Poland, closely resembles *Stenostomum gigerium* in most of the aspects, including a presence of the muscular gizzard composed of obliquely arranged muscle fibers. In the Polish worm, however, the gizzard is located immediately behind a short pharynx, while in the original description of *S. gigerium* the structure is described as interrupting the anterior portion of the intestine (Kepner and Carter 1931) in a manner characteristic for the other species in the genus *Myostenostomum* (e.g., Damborenea et al. 2011; Kostenko and Gaponova 2025; Luther 1960; Rogozin 1992; Tokinova and Berdnik 2017). Interestingly, Rogozin (1992), re-interpreted drawings of Kepner and Carter (1931), noting that the gizzard in *Myostenostomum gigerium*, unlike in other *Myostenostomum* species, is located immediately behind pharynx – a condition also observed in our worm. These historical disagreements regarding the position of the gizzard in relation to the pharynx are difficult to resolve. Unfortunately, the animals described in the literature have been documented exclusively through hand drawings, which are very prone to the authors’ personal interpretation. Until now, there is not a single attested photograph of *Stenostomum* (*Myostenostomum, Anokkostenostomum*) *gigerium* (*tauricum*).

Taking into account aforementioned taxonomic controversies surrounding *S. gigerium*, we decided to cautiously identify collected animals as *Stenostomum* cf. *gigerium*; a fourth report for this species from Poland (Kolasa 1971; Kolasa 1973; Kolasa 1977). Furthermore, we are also first to provide sequences of selected molecular markers for a putative representative of the genus *Myostenostomum*. To our surprise, the sequences of *S.* cf. *gigerium* show high similarity (100% for *28S*, *18S* and *COI* and 94% for *ITS 5.8*) to those of *S. hebuktense* described from Sweden. According to the description provided by Larsson and Willems (2010)., *S. hebuktense* possesses conspicuous pharynx equipped with well-developed musculature, but lacks a gizzard or similar structure. Unfortunately, the microphotographs of the live worms in the original publication are of insufficient quality to assess the detailed muscle arrangement and compare it with *S.* cf. *gigerium*. However, the longitudinal sections presented in the species diagnosis (Fig. 4F in Larsson and Willems (2010) show that radial muscles typical of the stenostomid pharynges, are restricted to the first half of the putative pharyngeal region. Further posteriorly, the lumen of the digestive system becomes constricted and numerous medio-lateral muscles fibers are inserted between the lumen and the body wall, resembling conditions observed in *S.* cf. *gigerium*.

By combining molecular and morphological evidence, we propose *S. hebuktense* as a younger synonym of *S. gigerium*. However, the relationship between *S. gigerium* and other species traditionally classified within *Myostenostomum* remains unresolved and requires further molecular data; particularly from species with a gizzard clearly separated from the pharynx, as expected for more typical representatives of *Myostenostomum*. An interesting question in this context is whether the gizzard of *S. gigerium* (*tauricum*, *hebuktense*) is homologous to that of typical *Myostenostomum* species, or whether these structures evolved convergently.

### Diversity of Catenulida in Poland

Until now all reports of Catenulida from Poland in its modern-day borders are based on morphological identification, primarily from the studies of Kolasa (Kolasa 1971; Kolasa 1973; Kolasa 1977; Kolasa and Young 1974a; Kolasa and Young 1974b) and, to a lesser extent, Moraczewski (Moraczewski 1981; Moraczewski et al. 1977). Together, these authors reported 23 catenulid species (including two found exclusively in the tropical palmhouses and aquaria in Poznań), from both Stenostomidae and Catenulidae families (Tab. 3). Our study adds three species new to Poland (*Stenostomum handoelense*, *Stenostomum* cf. *simplex* and *Stenostomum virginianum*), bringing the number of Polish catenulid species recorded from natural habitats to 24; a figure comparable to the well-studied catenulid fauna of Sweden (Larsson and Willems 2010) or the United States (Nuttycombe 1956; Nuttycombe and Waters 1938). The discovery of three previously unrecorded species from just four new localities, highlights that the studies of catenulid diversity in Poland are still in their infancy.

**Table 3.**
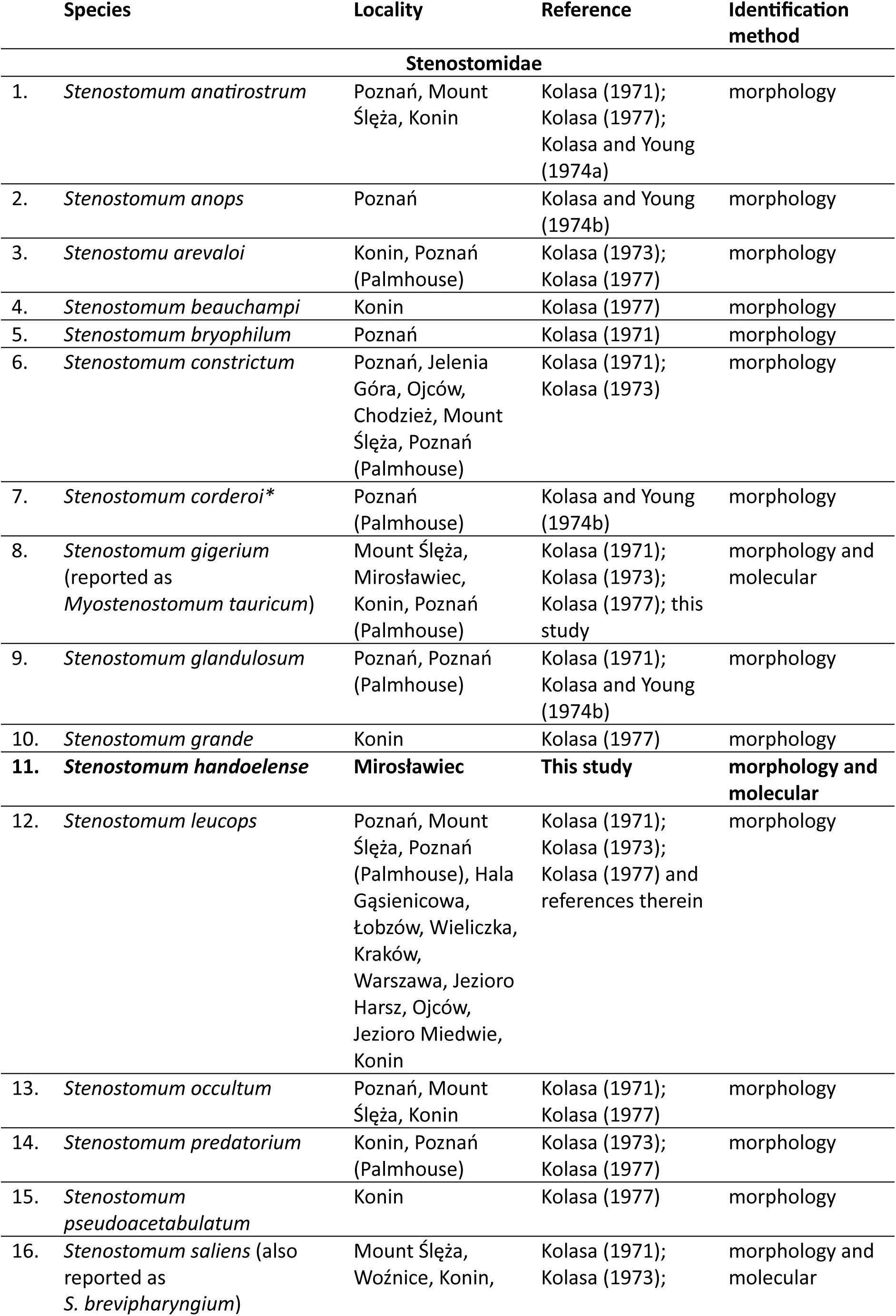
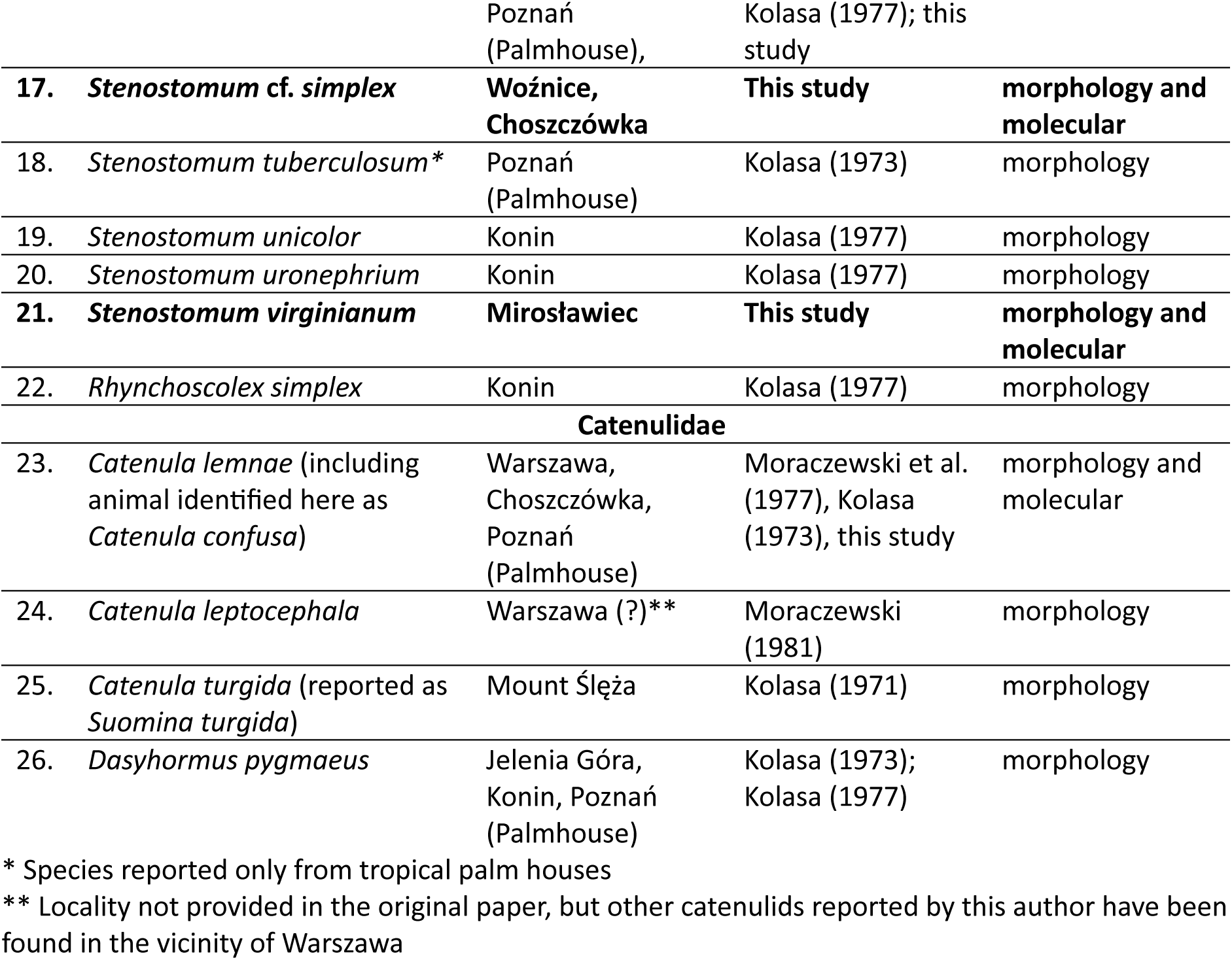
Catenulid species reported from Poland.

### Biogeography of Catenulida

The paucity of systematic surveys of catenulids and confusing taxonomy limit our understanding of their spatial distribution. Uneven sampling efforts, combined with uncertainties regarding the validity of some of the morphologically recognized species, prevent application of the rigorous, biogeographic analyses. To enable such studies, systematic sampling in additional regions, followed by sequencing and molecular identification, is essential. Nevertheless, even with the current limitations, some general observations on catenulid distribution patterns can already be made.

At least several catenulid species seem to have broad, transcontinental distribution. For instance, *S. saliens* has been reported from North America, South America, Japan and Europe (Kepner and Carter 1931; Kolasa 1971; Kolasa 1973; Kolasa 1977; Noreña et al. 2005; Nuttycombe and Waters 1938; Yamazaki et al. 2012, this study), and our molecular study confirms that the specimens from Japan, Germany and Poland are indeed conspecific. Interestingly, of the two Japanese individuals included in our analysis, one was placed closer to the European individuals, suggesting greater genetic diversity in Japan than in Europe. Similarly, one of the clades within *S. leucops* species complex groups animals from Europe, Asia, and Southern America that likely represent a single species.

We also provide the first report of *S. handolense* from Poland; a species previously known only from Sweden (Larsson and Willems 2010). However, in case of this species, and also, to a lesser extent, in *S. gigerium* (*S. heebuktense*), the individuals from Poland were molecularly quite well separated from the Swedish ones, suggesting reduced gene flow between Scandinavian and Central European populations. In contrast, we did not observe a similar pattern for *Catenula lemnae*, for which individuals from France, Germany and Poland were nested among animals derived from Swedish populations. These differences may reflect differences in dispersal capabilities or frequency of sexual reproduction, that could influence gene flow between individuals from the same location.

Overall, catenulids appear to exhibit low levels of endemism, with several species displaying broad, if not cosmopolitan, geographical range. Such distribution hints towards excellent dispersal capacities of catenulids, that has been also observed for several other microscopic invertebrates (e.g., Artois 2011; Cerca et al. 2018; Sato et al. 2025; Tessens et al. 2021). In the case of catenulids, the potential for dispersal might be further enhanced by asexual reproduction that allows even a single individual to establish a viable population.

### Evolution of morphological, developmental and ecological characters

Eight of the sixteen characters analyzed in our study revealed phylogenetically informative patterns, highlighting potential morphological synapomorphies for major catenulid clades (see taxonomic remarks in the Discussion). In accordance with the results of Janssen et al. (2015), we found evidence for independent evolution of paratomy in Microstomidae and Catenulida. Moreover, within catenulids, the paratomy seems to be lost independently in Retronectidae and Paracatenulidae. Finally, we also tested the hypothesis proposed by Leisch et al. (2011), which suggested that marine Retronectidae *sensu lato* (including *Paracatenula*) are derived from freshwater ancestors. This evolutionary scenario could explain the apparent reduction of protonephridial organs in those worms (Sterrer and Rieger 1974). Our reconstruction supports this hypothesis, further indicating that transition into marine habitats occurred indepdentnly in Retronectidae *sensu stricto* and Paracatenulidae (Fig. 6).

The evolution of eight remaining characters was more difficult to reconstruct. Four of them – excretophores, epidermal inclusions, long stiff cilia across the body, and ability to form chains – appear to be highly evolutionarily labile. In contrast, the ambiguous reconstruction for upside-down swimming likely results from the missing behavioral data for many species in our dataset. This behavior has only been recorded in members of the Clade 4 and is present in all scored representatives of this clade. We suspect that, with additional data, the upside-down swimming would be reconstructed as ancestral characteristic of Clade 4. As this character is easy to observe and appears phylogenetically restricted to Clade 4, it could be of a great use in identification keys or subgeneric taxonomy of *Stenostomum*. The three remaining characters – muscular gizzard, protonephridium and eyespots – exhibited little to no variation across the analyzed species.

## 5. Conclusions

Integrative taxonomy, that combines morphological and molecular data, is a powerful tool for studying catenulid diversity. Multi-marker phylogenies not only allow for more precise species identification, but also create framework for biogeographic, morphological, developmental, and ecological research. Here, we used such integrative approach to delimit the major clades within Stenostomidae, the largest family of catenulids, and to trace the evolution of taxonomically important characters, identifying potential synapomorphies for key lineages.

Although an increasing number of catenulid species are being sequenced, considerable gaps remain in the molecular coverage of their diversity. Notably, most of the catenulid sequences come from Europe, which remains the best studied region using molecular tools. Future sequencing efforts should prioritize regions with high catenulid diversity that have been studied almost exclusively with morphological methods (e.g., South and North America), as well as regions that remain largely unexplored (e.g., South-East Asia, Africa).

## 6. Supporting Information

- **FileS1_alignment.fasta** Nucleotide alignment used in phylogenetic analysis
- **FileS2_matrix.txt** Matrix with characters used in ancestral state reconstructions
- **FileS3_MLtree.tre** Maximum likelihood tree with all bootstrap values
- **FileS4_18S_trim_FastTree.tre** Approximate maximum-likelihood tree for *18S* with SH-like branch support
- **FileS5_28S_trim_FastTree.tre** Approximate maximum-likelihood tree for *28S* with SH-like branch support
- **FileS6_COI_trim_FastTree.tre** Approximate maximum-likelihood tree for *COI* with SH-like branch support
- **FileS7_ITS_trim_FastTree.tre** Approximate maximum-likelihood tree for *ITS-5.8* with SH-like branch support
- **TableS1.xlsx** Accession numbers of nucleotide sequences used in the analysis
- **TableS2.docx** Results of model fitting for the ER and ARD models applied to the traits analyzed in this study
- **Fig. S1.** The ancestral state reconstruction of the characters with weak phylogenetic signal.

## 7. Data Availability Statement

The molecular data underlying this article are available in the GenBank Nucleotide Database under accession numbers PQ887791–PQ887810, PQ895509–PQ895524, PV019496–PV019509 and PV403740–PV403751. The remaining data analyzed during this study are included in this article and its supplementary materials.

## Acknowledgments

We would like to thank our collaborators who provided worms from various localities across Western Europe: Nicolas Bekkouche from the Sorbonne University (animals from France), Chris Laumer from the Natural History Museum London (animals from the UK) and Matteo Vecchi from The Institute of Systematics and Evolution of Animals of the Polish Academy of Sciences (animals from Italy). We are also very grateful to Bożena Zakryś from the University of Warsaw, who shared some of her freshwater samples and advised us on the DNA extraction method. Alexander Kostenko from the Institute for Evolutionary Ecology of The National Academy of Sciences of Ukraine is greatly acknowledged for insightful discussion on the taxonomy of the genus *Myostenostomum*.

## 9. Authors’ contributions

KT collected, photographed and cultured the animals, performed molecular laboratory work, processed the sequences, compiled morphological matrix, prepared figures and drafted parts of the manuscript. JB perform phylogenetic analyses, prepared figures and drafted parts of the manuscript. LG designed the study, acquired funding, collected samples, identified and photographed the animals, perform phylogenetic analyses, compiled morphological matrix, prepared figures and drafted the manuscript. All authors read and accepted the final version of the manuscript.

